# Structure and Epitope of a Neutralizing Monoclonal Antibody that Targets the Stem Helix of β Coronaviruses

**DOI:** 10.1101/2022.09.14.507947

**Authors:** Ashlesha Deshpande, Nobert Schormann, Mike S. Piepenbrink, Luis Martinez Sobrido, James J. Kobie, Mark R. Walter

## Abstract

Monoclonal antibodies (MAbs) that retain neutralizing activity against distinct coronavirus (CoV) lineages and variants of concern (VoC) must be developed to protect against future pandemics. These broadly neutralizing MAbs (BNMAbs) may be used as therapeutics and/or to assist in the rational design of vaccines that induce BNMAbs. 1249A8 is a BNMAb that targets the stem helix (SH) region of CoV spike (S) protein and neutralizes Severe Acute Respiratory Syndrome Coronavirus 2 (SARS-CoV-2) original strain, delta, and omicron VoC, Severe Acute Respiratory Syndrome CoV (SARS-CoV) and Middle East Respiratory Syndrome CoV (MERS-CoV). To understand its mechanism of action, the crystal structure of 1249A8 bound to a MERS-CoV SH peptide was determined at 2.1Å resolution. BNMAb 1249A8 mimics the SARS-CoV-2 S loop residues 743-749, which interact with the C-terminal end of the SH helix in the S postfusion conformation. The crystal structure shows that BNMAb 1249A8 disrupts SH secondary structure and packing rearrangements required for CoV S to adopt its prefusion conformation that mediates membrane fusion and ultimately infection. The mechanisms regulating BNMAb 1249A8 CoV S specificity are also defined. This study provides novel insights into the neutralization mechanisms of SH-targeting CoV BNMAbs that may inform vaccine development and the design of optimal BNMAb therapeutics.

## Introduction

Coronaviruses (CoV) infect animals and humans causing respiratory disease with varied severity. Endemic human CoV, such as OC43 and HKU1, often cause cold-like symptoms [1–3]. In contrast, three CoV have crossed the animal-human species barrier, generating highly pathogenic viruses responsible for large numbers of human fatalities. Severe Acute Respiratory Syndrome CoV (SARS-CoV) emerged in China in 2002 and was responsible for 774 human fatalities [4, 5]. In 2012, the Middle Eastern Respiratory Syndrome CoV (MERS-CoV) emerged on the Saudi Arabian Peninsula, spread to 27 countries, and caused 791 human fatalities as of 2018 [6]. MERS-CoV continues to circulate with the latest fatality reported in May, 2022 [7]. In addition to these epidemics, Severe Acute Respiratory Syndrome CoV-2 (SARS-CoV-2) emerged in the city of Wuhan, China in 2019 and has been responsible for the Coronavirus Disease 2019 (COVID-19) pandemic that, to date, has claimed the lives of over six million people worldwide [8, 9].

Significant genome diversity has been observed in beta-coronaviruses (β-CoV) that infect humans. OC43 and HKU1 belong to lineage A, SARS-CoV and SARS-CoV-2 to lineage B, and MERS-CoV to lineage C β-CoV [10, 11]. Sequence analyses strongly suggest SARS-CoV, SARS-CoV-2, and MERS-CoV originated in bats [12–15], while civets (SARS-CoV and SARS-CoV-2) and dromedary camels (MERS-CoV) are intermediary hosts that transmit the viruses to humans [15, 16]. CoV with sequences related to SARS-CoV (e.g. SARSr-CoV) have been found in bats located across Asia [17]. More recently, SARSr-CoV sequences, containing all the genetic elements required to form SARS-CoV, were isolated from bats found in a single cave in Yunnan province, China [18]. Identifying one location containing extensive SARS-CoV genetic diversity, combined with the ability of CoV to undergo recombination [19, 20], suggests hotspots exist that are poised for generating future zoonotic CoV [16]. In summary, all evidence suggests CoV spillovers into humans will occur in the future [15, 21, 22] and the development of broadly neutralizing antibodies against these CoV, and vaccines that induce such antibodies, will be important for future pandemic preparedness.

The CoV spike (S) protein is responsible for viral entry into host cells. S adopts a homotrimer where each subunit consists of two domains, S1 and S2 [23–25]. Viral attachment to host cells is mediated by S1, through the receptor binding domain (RBD), while the S2 domain mediates membrane fusion. On the viral surface, S adopts a mushroom shaped pre-fusion (PreF) conformation, with S1 RBDs located furthest away from the viral membrane [26, 27]. Host receptor binding requires the RBDs to undergo conformational changes that, along with protease cleavage events, allow the meta-stable S2 domain to refold into a low energy rod-shaped coiled-coil structure, called the post-fusion (PostF) S conformation [26]. Structures of preF and postF S provide the beginning and end conformations of S that lead to membrane fusion [26].

Sequence and structural variation among CoV lineages and variants is not equivalent throughout the S polypeptide. S1 exhibits high sequence, structure, and functional diversity (e.g. required to bind distinct host receptors), while the S2 domain is more conserved. Variability in S1 alters cell tropism and/or receptor binding affinity to improve viral fitness. For example, SARS-CoV-2 S1 has rapidly acquired amino acid changes, resulting in viruses defined as variants of concern (VoC), that have altered pathogenesis and transmission properties, relative to the initially isolated Wuhan virus [28]. S1 is also under intense host immune pressure, where studies estimate ~75%-90% of the humoral immune response is targeted to S1 [29, 30]. As a result, most neutralizing monoclonal antibodies (MAbs) and vaccines that target SARS-CoV-2 S1 N-terminal and RBD regions, exhibit reduced neutralizing potency to evolving VoC and little to no efficacy against different CoV lineages, including SARS-CoV and MERS-CoV [31, 32].

In contrast to S1, S2 exhibits higher sequence and structure conservation across β-CoV lineages and has not undergone significant antigenic drift between SARS-CoV-2 VoC. This suggests epitopes within the S2 domain may be optimal targets for the development of BNMAbs against diverse present-day and future emerging CoV. To address this hypothesis, we previously reported a series of human S2-targeting neutralizing MAbs [33]. One MAb, 1249A8, was found to neutralize SARS-CoV-2 and VoC (delta and omicron), SARS-CoV, and MERS-CoV [33]. BNMAb 1249A8 targets the stem helix (SH) region of β-CoV S protein and is the only human SH MAb [34–37] reported to neutralize MERS-CoV [33].

To understand the structural basis for 1249A8’s broad specificity and potency, the crystal structure of BNMAb 1249A8 Fab, in complex with a stem helix peptide (SHp) from MERS-CoV S, was determined at 2.1Å resolution. The structure identifies the unique specificity determinants of the 1249A8-MERS-CoV SH interaction required for broad CoV binding and suggests 1249A8 inhibits membrane fusion by disrupting SH secondary structure rearrangements and sterically occluding SH-CH/HR1 interactions that are required for preF S to transition to its postF conformation.

## Results

### Structure of the 1249A8/MERS-CoV SH complex

The crystal structure of 1249A8 Fab bound to a MERS-CoV S SHp, residues 1,223 to 1,245, was solved at 2.1Å resolution (**Fig. 1, Supp. Table 1**). Of the 23 residues in the peptide, electron density was observed for 12 SHp residues (1,230-1,241) that form an amphipathic α-helix that binds in a groove between heavy and light chain CDRs of 1249A8. The MERS-CoV SHp sequence corresponds to SARS-CoV-2 residues 1,147-1,158. Three structures of human NAbs binding to the SARS-CoV-2 SH have been determined (**Fig. 1**, and refs. [34–39]). Fab-SHp structure comparisons reveal 1249A8, S2P6 and CC40.8 exhibit similar, but distinct, epitopes, defined as class-1 (C1) NAbs, that bind predominantly to the hydrophobic surface of the SH. Based on the cryo-EM structure of SARS-CoV-2 S, derived from the native S sequence [26], the C1 NAb epitope is buried in the center of the trimeric SARS-CoV-2 pre-fusion S coiled-coil (CC) domain (**Fig. 1b**). In contrast, CV3-25, defined as a class-2 (C2) NAb, targets exposed hydrophilic residues on SH and exhibits a distinct binding orientation, relative to the other NAbs (**Fig. 1b**). The C1 NAbs bind to SH using CDRs from their H and L chains, while the C2 CV3-25 binds almost exclusively (95%) through H chain CDR residues.

**Figure 1.**
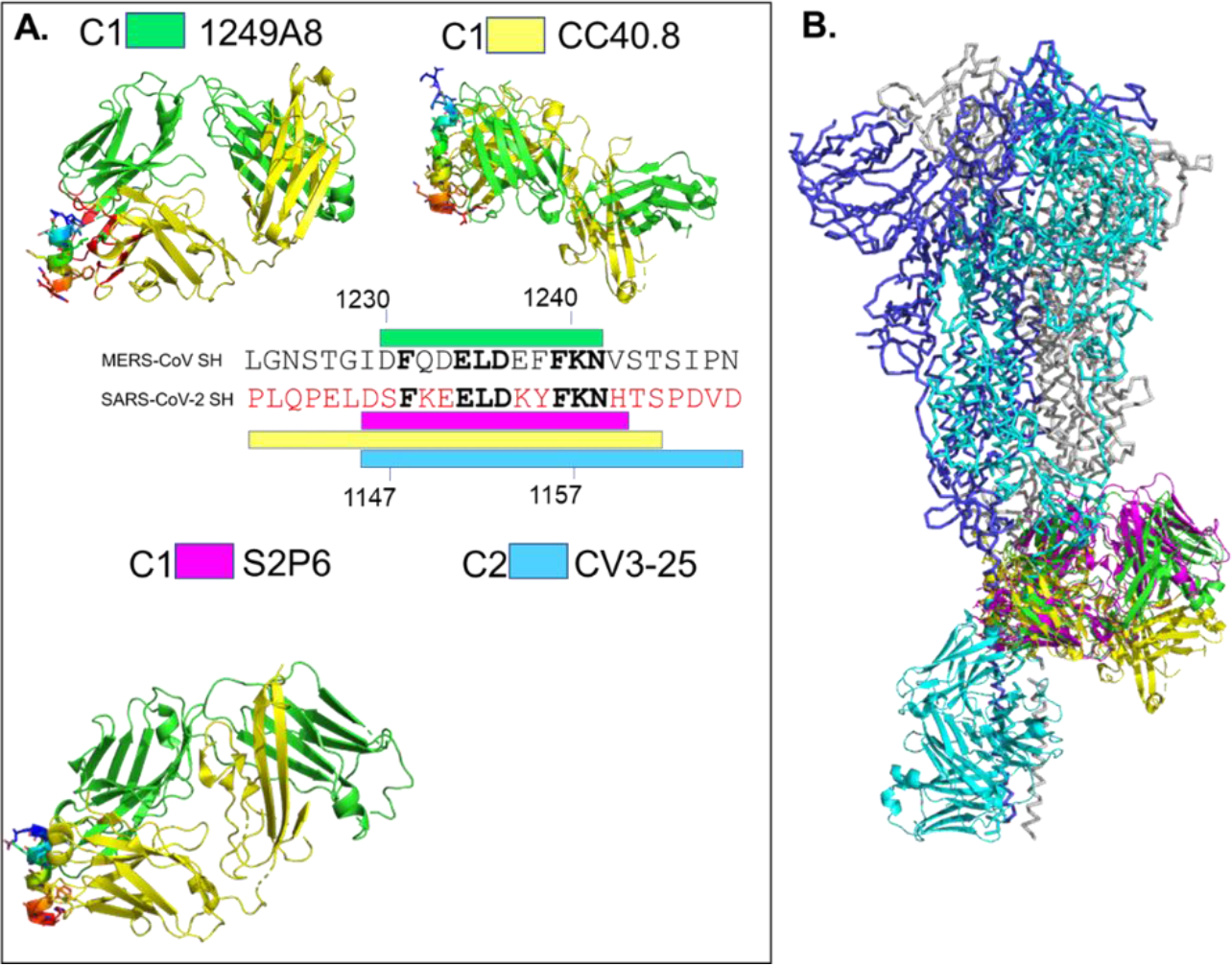
Structure of 1249A8/MERS-CoV SHp complex and related SH-targeting NMAbs. (a) Ribbon diagram of the 1249A8/MERS-CoV SHp complex and other human NMAb/SHp complexes. 1249A8 heavy and light chains are colored yellow and green, respectively. The MERS-CoV SHp is rainbow colored from the N-terminus (blue) to the C-terminus (red). Other NAb-SHp structures are colored as described for 1249A8. Labels for each NMAb-SH complex contain the designation of C1 (class-1) or C2 (class-2) to distinguish their epitopes. The linear SH epitope for each NMAb is shown on the MERS-CoV (black) and SARS-CoV-2 (red) sequences as a colored bar. Conserved residues in the linear sequence are bolded. The invariant core-SH region is highlighted by the black bar. (b) NMAb binding epitopes are superimposed on a model of the SARS-CoV-2 pre-fusion S trimer (6xr8) and the extended C-terminal helix residues 1,171-1,203 (pdbid 6lvn). NMAbs are color coded as shown in (a).

Superposition of 1249A8, and other C1 NAbs, onto the SH region of the preF S (pdbid 6xra) results in numerous steric clashes between S and Fab, suggesting that distortion / disruption of the trimeric CC region of S is required for NAb binding (**Fig. 1b**). Cryo-EM of the S2P6-SARS-CoV-2 S complex, a C1 SH-targeting NAb, clearly supports this hypothesis [34]. In contrast, the surface exposed binding site of CV3-25 would induce limited distortion of the CC region (**Fig. 1b** [37]). Structures of S from MERS-CoV, HKU1, SARS-CoV and SARS-CoV-2 (WA-1 strain) show the SH/CC region is disordered in 2P proline-stabilized S structures [23–25, 40, 41]. Furthermore, tomograms of SARS-CoV-2 particles reveal the SH / CC region of S undergoes a variety of bending motions that are presumably important in the membrane fusion process [42, 43]. Together, these data suggest both C1 and C2 human NAb SH epitopes are at least temporarily accessible in the preF S conformation.

### 1249A8 exhibits higher affinity for SH peptide than prefusion S

We previously defined the affinity of 1249A8, and other SH-targeting NAbs, for the SARS-CoV-2 PreF S [33]. Here, we defined the affinity of 1249A8 to a SARS-CoV-2 SH peptide (residues 1,131-1,171) to determine the role of S structure on 1249A8 binding (**Fig. 2**). 1249A8 bound preF S ~8-fold weaker (*K*D = 3.29nM) than the SH peptide (*K*D = 0.42nM). 1249A8 exhibited the highest affinity for SH peptide, when compared to S2P6, CV3-25, and CC40.8, although the differences were small (~2-fold), relative to NAb affinity differences observed when binding to S monomer (~10-fold, **Table 1**). The higher binding affinity of the NAbs to the SH peptide may be at least partially due to avidity, as the SH peptide was expressed as an FC fusion protein. In addition to affinity, SH epitope accessibility in S trimers could influence NAb binding and function. To address this possibility, C1 and C2 NAb binding levels to various S proteins were determined (**Fig. 2**). Proteins evaluated in the study included the SH peptide, trimeric SARS-CoV-2 omicron VoC, expressed as a 2P or 6P stabilized S, and trimeric MERS-CoV S. Different omicron S proteins were evaluated since it has been reported that the 6P variant is more stable than the 2P form of S [44]. The resulting surface plasmon resonance (SPR) data was consistent with previously determined affinity and specificity data of 1249A8, relative to other C1 and C2 NAbs. Specifically, 1249A8 exhibited the highest binding levels to the SH peptide, SARS-CoV-2 omicron-2P S, and MERS-CoV S. However, C2 NAb CV3-25 exhibited the highest binding to level to SARS-CoV-2-6P, while the three C1 NAbs all exhibited lower binding levels that were similar to each other. This data is consistent with the surface accessibility of the CV3-25 epitope, relative to the C1 NAbs (**Fig. 1b**).

**Figure 2.**
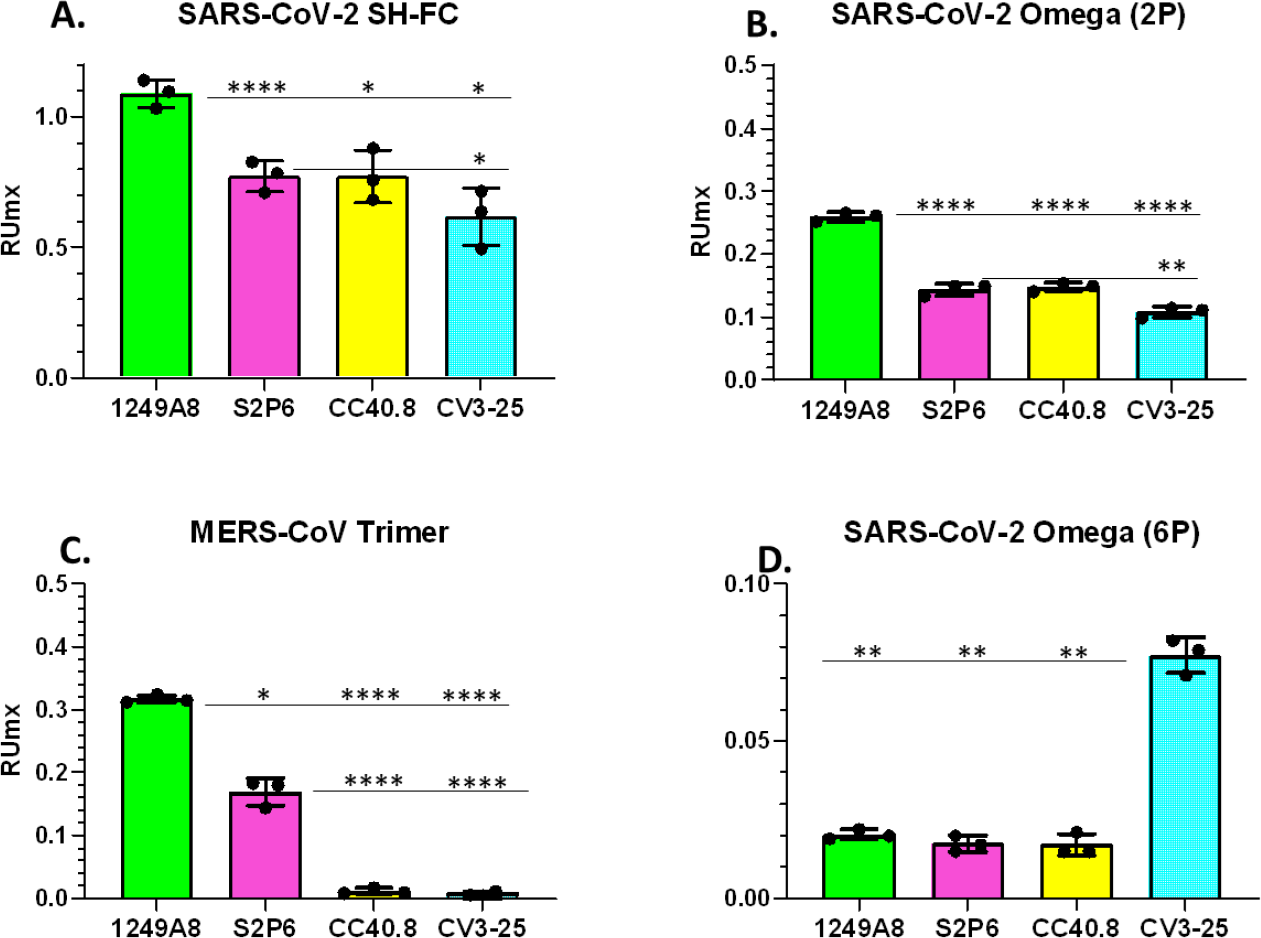
NMAb binding to SH peptides and SH within CoV S proteins. Fractional binding of SH peptide (A) and CoV S proteins: SARS-CoV-2-2P omega B.1.1.529 S (B) MERS-CoV S (C) and SARS-CoV-2-6P omega B.1.1.529 S (D).

**Table 1.**
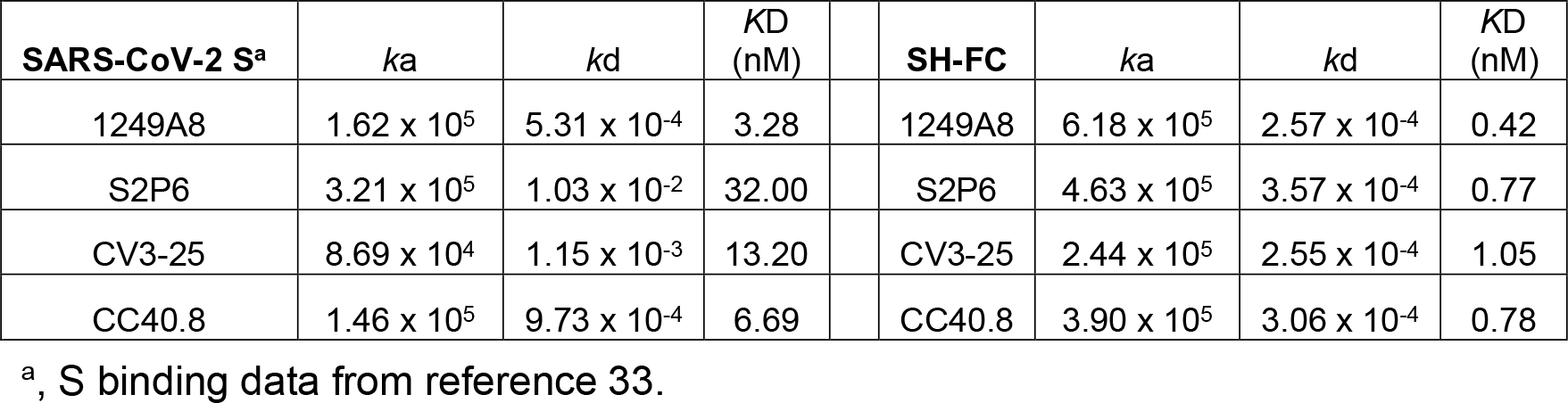
NAb Binding Parameters.

| SARS-CoV-2 S<sup>a</sup> | $k_a$ | $k_d$ | $K_D$ (nM) | SH-FC | $k_a$ | $k_d$ | $K_D$ (nM) |
| --- | --- | --- | --- | --- | --- | --- | --- |
| 1249A8 | $1.62 \times 10^5$ | $5.31 \times 10^{-4}$ | 3.28 | 1249A8 | $6.18 \times 10^5$ | $2.57 \times 10^{-4}$ | 0.42 |
| S2P6 | $3.21 \times 10^5$ | $1.03 \times 10^{-2}$ | 32.00 | S2P6 | $4.63 \times 10^5$ | $3.57 \times 10^{-4}$ | 0.77 |
| CV3-25 | $8.69 \times 10^4$ | $1.15 \times 10^{-3}$ | 13.20 | CV3-25 | $2.44 \times 10^5$ | $2.55 \times 10^{-4}$ | 1.05 |
| CC40.8 | $1.46 \times 10^5$ | $9.73 \times 10^{-4}$ | 6.69 | CC40.8 | $3.90 \times 10^5$ | $3.06 \times 10^{-4}$ | 0.78 |

<sup>a</sup>, S binding data from reference 33.

### 1249A8/MERS-CoV SHp Interface

The MERS SHp α-helix buries 614Å^2^ of accessible surface area into 1249A8, which is distributed between heavy (368 Å^2^) and light (246 Å^2^) chain CDRs. Hydrophobic residues F1231 L1235, and F1239 bury the greatest amount of surface area into 1249A8 (**Fig. 3, supplemental Fig. 1**).

**Figure 3.**
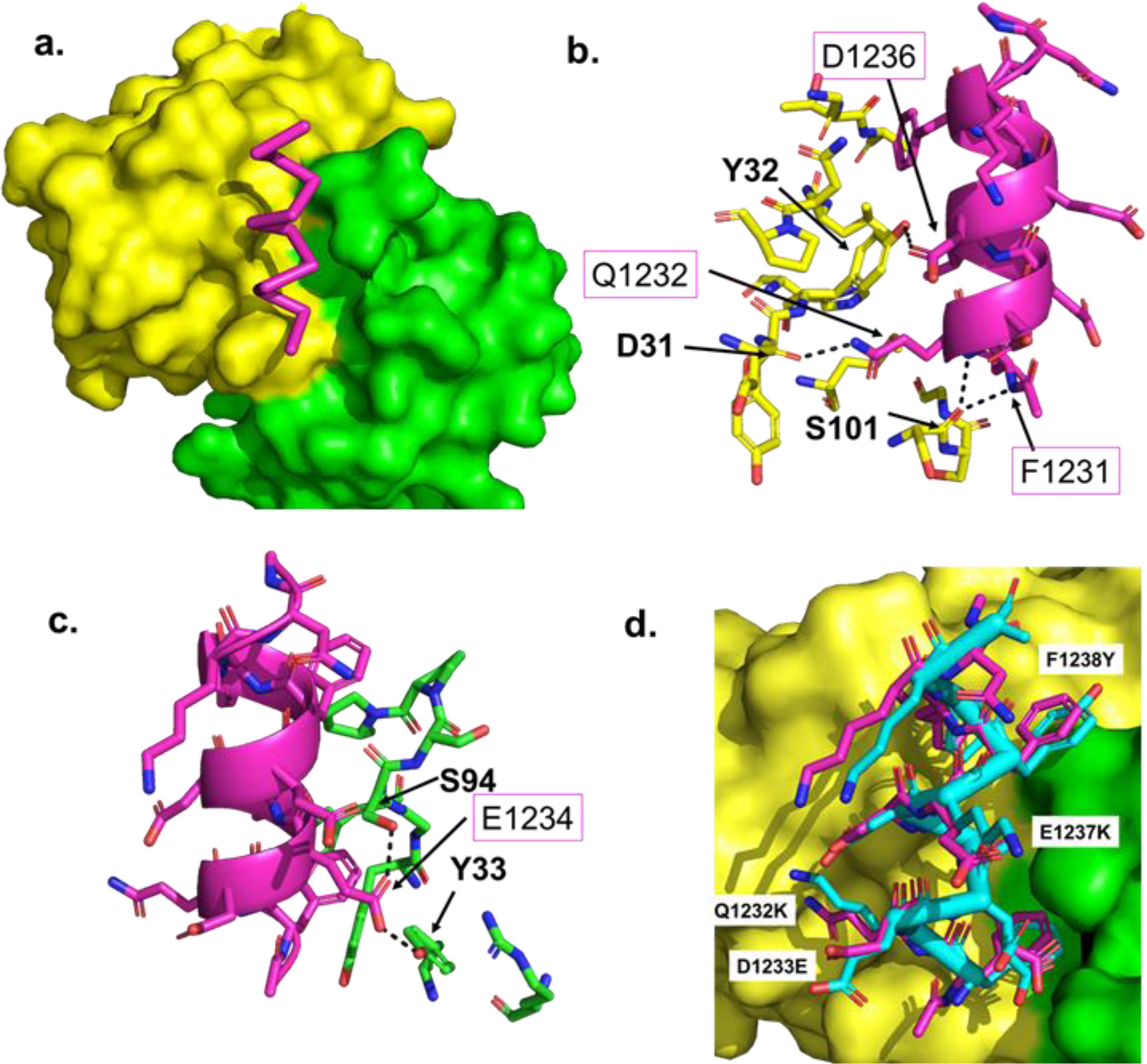
Details of the 1249A8/MERS-CoV SHp epitope. **(a)** Surface representation of 1249A8 with heavy and light chains colored as in Figure 1, with the MERS-CoV SHp shown in magenta. Optimized views of the MERS-CoV SHp interacting with the 1249A8 heavy **(b)** and light **(c)** chains. **(d)** Superposition of SARS-CoV-2 SHp (7rnj) onto the MERS-CoV SHp. The superimposed SHps are shown with the 1249A8 Fab surface showing that both SHps fit in the 1249A8 binding pocket without steric clashes.

Other SHp residues with hydrophobic and hydrophilic chemistries also bury significant amounts of surface area into 1249A8. However, these residues remain partially accessible to the solvent to accommodate alternative amino acid residues found in other CoV SH regions. A total of six hydrogen bonds are made between 1249A8 CDRs and the MERS-CoV SHp (**Figs. 3b, c, Table 2**). Four are with heavy chain residues and two are with the light chain. Five of the six hydrogen bonds are formed with four N-terminal residues (1,231-1,234) of the epitope. Thus, while the 1249A8 epitope consists of residues 1,231-1,239, critical 1249A8 binding contacts are concentrated into the four-residue segment, residues 1,231-1,234. Two of the six hydrogen bonds are formed between the carbonyl oxygen of CDRH3 S101 and the backbone nitrogen atoms of SHp F1231 and Q1232. To form these hydrogen bonds, CDRH3 is positioned beneath the N-terminal end of the helix, effectively “capping” the SHp. While less pronounced, other C1 NAbs share a similar binding mode, suggesting C1 NAbs require the full length preF α-helical SH to unwind and/or bend to allow NAb binding.

**Table 2.** Hydrogen bonds in the 1249A8 / MERS-CoV SHp complex.

| Chain | # | MERS SHp Atom | Dist. [Å] | 1249A8 Atom |
| --- | --- | --- | --- | --- |
| HC | 1 | PHE1231[ N ] | 2.80 | SER 101[ O ] |
| HC | 2 | GLN1232[ N ] | 2.86 | SER 101[ O ] |
| HC | 3 | GLN1232[ NE2] | 2.71 | ASP 31[ O ] |
| HC | 4 | ASP1236[ OD1] | 2.50 | TYR 33[ OH ] |
| LC | 5 | GLU1234[ OE1] | 2.45 | SER 94[ OG ] |
| LC | 6 | GLU1234[ OE2] | 2.24 | TYR 33[ OH ] |

### Molecular Basis for 1249A8’s Broad Specificity

While all reported SH-targeting human NAbs bind to the SARS-CoV-2 SH, 1249A8 is the only NAb that also exhibits high affinity binding (*K*D = 0.58nM) to the MERS-CoV SH and neutralizing activity against MERS-CoV [33]. The six amino acid differences between the MERS-CoV and SARS-CoV-2 SH epitopes have limited impact on 1249A8 interactions because they are either fully (D1229S, D1233E, E1237K, V1242H), or partially (Q1232K and F1238Y), accessible to solvent when bound to 1249A8. The Q^MERS-CoV^1232K^SARS-CoV-2^ mutation is the only residue substitution that alters the hydrogen bonding found in the 1249A8-MERS-CoV SHp complex (**Fig. 3d**). 1249A8 CDRH1 accommodates both Q and K sidechains by forming a hydrogen bond between Q1232 and the mainchain carbonyl of D31, while SARS-CoV-2 K1149 is positioned to interact with the negatively charged D31 sidechain (**Fig. 3b**). In both cases, the common aliphatic regions of MERS-CoV Q1232 and SARS-CoV-2 K1149 sidechains bury significant amounts of surface area into the 1249A8 interface to maintain high affinity binding. In contrast to 1249A8, other C1 NAbs exhibit weak (S2P6, *K*D 8.5nM) or essentially no affinity (CV3-25 and CC40.8) for the MERS-CoV SH. Analysis of modeled S2P6/MERS-CoV SH and CC40.8/MERS-CoV SH complexes show replacement of K1149 with Q induces steric clashes within the S2P6 and CC40.8 binding sites, consistent with their low or absent affinity for MERS-CoV SH (**Fig. 3d**). Q1232 cannot be responsible for CV3-25’s inability to bind to MERS-CoV SH, since the residue is not part of the binding epitope. However, 12 additional SARS-CoV-2 SH residues that form the CV3-25 binding epitope are different in the MERS-CoV SH (**Fig. 1**) and all could impact CV3-25/MERS-CoV SH interactions.

### 1248A8 CDRH3 mimics the SARS-CoV-2 S 3-helix motif

Based on the static preF SARS-CoV-2 S structure, optimal 1249A8 binding requires dissociation of trimeric S to remove Fab-S steric clashes and expose the 1249A8 epitope (**Fig. 1b**). In addition to trimer dissociation, 1249A8 cannot bind efficiently to the full-length α-helical SH (residues 1,141-1,161) due to steric clashes with CDRH3, which binds across the N-terminal end of SH residue F1148 (**Fig. 3b**). Thus, optimal 1249A8 binding requires the preF SH to unwind. SH unwinding occurs naturally as S transitions from its preF to postF conformation (**Fig. 4a**), changing from a 21-residue α-helix in preF S to a 7-residue core α-helix (core-SH, residues F1148-K1154) in the postF S (**Figs. 1, 4b**). The core-SH is structurally conserved in preF and postF S conformations, exhibits high sequence conservation across CoV lineages, and contains the essential five 1249A8 binding residues required to form all six hydrogen bonds with 1249A8. This suggests 1249A8 is optimized to bind the postF SH. Superposition of 1249A8-SHp complex onto the core-SH in the postF S (6xra) reveals CDRH3 mimics the loop structure (residues 743-749, CGDSTEC) found in the 3-helix region (residues 737-769) of the postF S, where D745 caps the N-termini of the SH (**Fig. 4c**).

**Figure 4.**
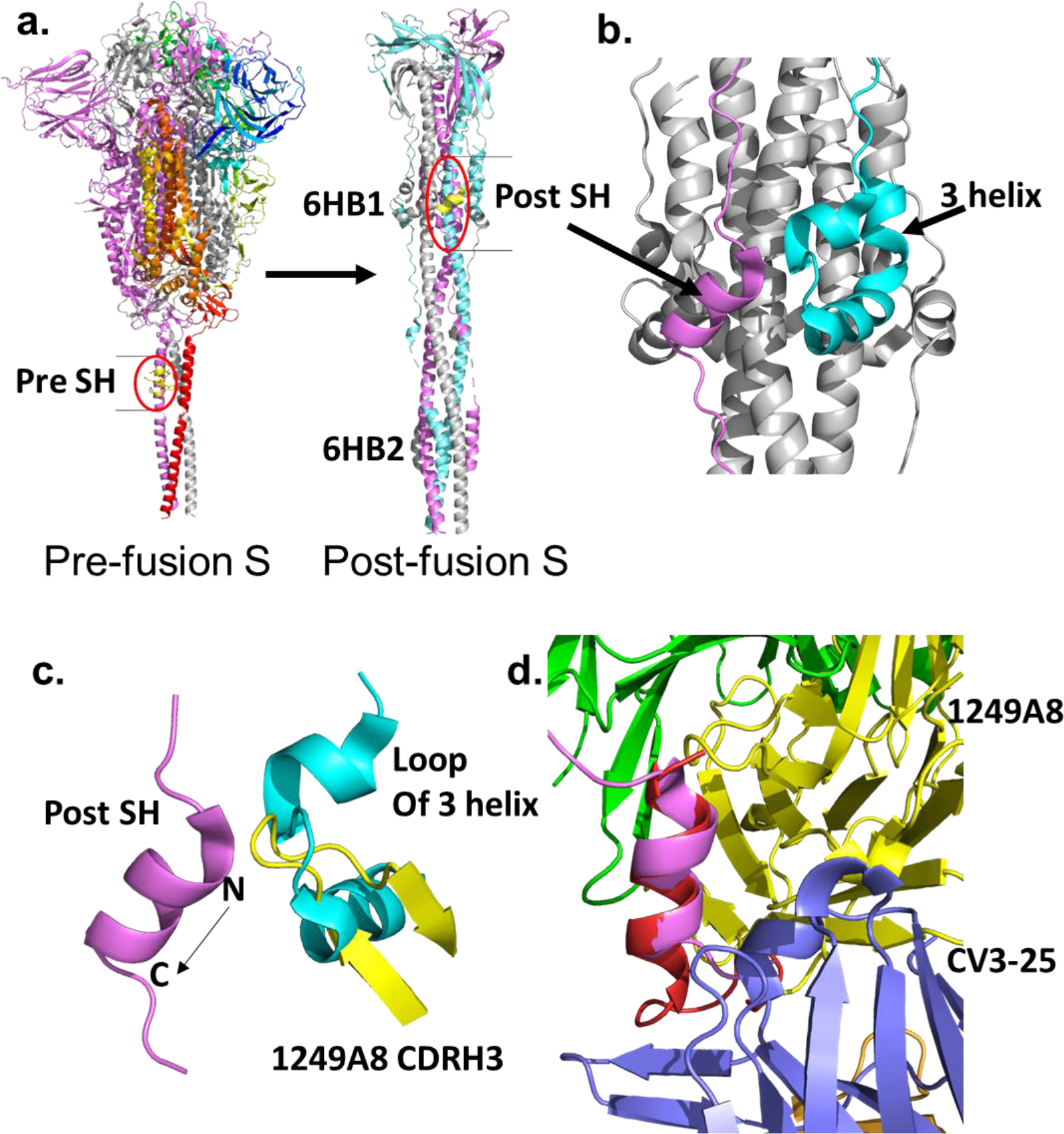
SARS-CoV-2 pre- and post-fusion structures and mimicry by 1249A8. **(a)** Location of the SHps in the structures of the SARS-CoV-2 pre-fusion (6rx8+6lvn) and post-fusion (6xra) S structures. The location of 6HB1 and 6HB2 are shown on the post-fusion S. **(b)** Packing of the SHp (pink) with the 3-helix region (cyan) against the CH region. **(c)** 1249A8 (yellow) mimics the SARS-CoV-2 S 3-helix region loop (residues 743-749), which caps the N-terminal end of the post-SH core helix. **(d)** Distinct SH binding epitopes that target the N-terminal of end of SH (1249A8) and the C-terminal end of SH (CV3-25).

This data suggests 1249A8-mediated neutralization occurs by blocking 6HB1 assembly (**Fig. 4a**), and ultimately disrupting formation of the postF S structure required for membrane fusion. All C1 NAbs bind the N-terminal end of the postF core-SH and are hypothesized to neutralize virus by sterically interfering with 6HB1 formation as described for 1249A8. In contrast, C2 CV3-25 binds to the C-terminal end of the postF core SH on the accessible surface of the helix (**Fig. 4d**). As a result, CV3-25 does not block the SH/3-helix interaction and makes only minor steric clashes to prevent core-SH packing with CH/HR1 to form the 6HB1. In comparison to C1 NAbs, CV3-25 would be predicted to have limited ability to disrupt the formation of the postF S, if steric disruption of 6HB1 formation was the only mechanism used by SH-targeting NAbs.

### 1249A8 disrupts post-fusion SH secondary structure

The SARS-CoV-2 SH makes an extensive transition from a preF 21-residue helix to a postF extended structure that retains a 7-residue α-helical core-SH, which changes the length of SH by 16Å (**Fig. 5**).

**Figure 5.**
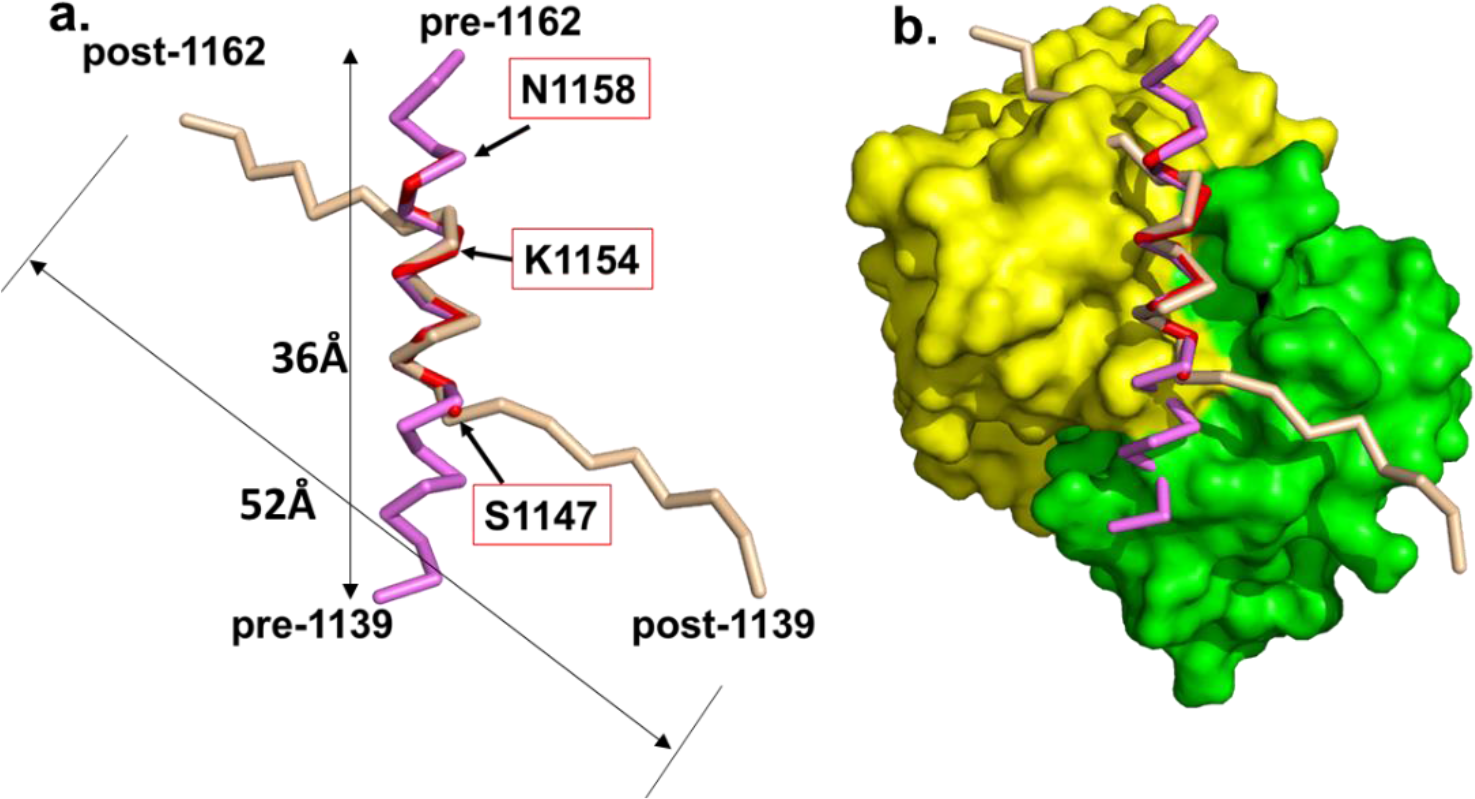
1249A8 locks SH in a pre-fusion α-helical conformation. **(a)** Ribbon diagram with pre- and post-fusion SH regions (residues 1139-1162) from pdbids 6xr8 and 6xra superimposed onto the 1249A8 binding epitope (SARS-CoV-2 SH residues 1147-1158, red). Boxed residues highlight the residues mimicking the post-fusion core α-helix (1147-1154) and the end of the epitope (1158). **(b)** Superposition of (a) is shown in the context of the 1249A8 Nab surface, which shows 1249A8 cannot bind to the full-length pre-fusion α-helix and how 1249A8 disrupts the secondary structure of the post-fusion SH, locking it into the pre-fusion conformation. 1249A8 binding disrupts this SH transition by locking the SH in a pre-fusion α-helical conformation.

1249A8 forms all hydrogen bonds with the naturally occurring postF core-SH (F1148-K1154). However, the remainder of the 1249A8 epitope (Y1155-N1159) retains an α-helical conformation observed in the preF SH, while the same residues in postF SH adopt an extended conformation required for proper postF S refolding (**Fig. 5b**). Thus, 1249A8 binding disrupts SH secondary structure transitions required for SH and HR2 regions to properly pack against HR1 (**Figs. 4,5**). Thus, 1249A8 is predicted to disrupt postF S 6HB1 and 6HB2 formation using steric and disrupted SH secondary structure mechanisms. All SH-targeting NAbs are expected to neutralize CoVs using similar mechanisms.

## DISCUSION

Given the likelihood of future pandemics, including those caused by CoV, there is a great need to develop BNMAbs against current and future CoV lineages and optimal vaccines that elicit them. 1249A8 is a human BNMAb that targets the SH region of S and neutralizes β-CoV from two lineages (B SARS-CoV-1 and SARS-CoV-2, including VoC, and C MERS-CoV), extending the breadth of protection realized with current human BNMAb therapies [33]. While 1249A8 is the only human SH-specific BNMAb that neutralizes MERS-CoV, murine-derived SH-targeting NAbs (1.6C7, 28D9, Fab22, B6) with activity against MERS-CoV have been isolated [45–47]. The murine NAbs were generated by immunizations with full-length preF S, or stabilized preF S2, and not SH peptides, further establishing the immunodominance of β-CoV SH region in mice and humans [48, 49]. Overall, the data suggests SH is an important target for the development of broad specificity NAbs and a target for vaccine optimization.

It has been difficult to rationalize the proposed neutralization mechanisms of SH-targeting NAbs with their binding and structural attributes. For example, structural studies demonstrate the murine NAbs, Fab22 and B6, target a similar buried hydrophobic MER-CoV SH epitope as 1249A8 (e.g. C1 NAbs). However, Fab22 and B6 neutralize MERS-CoV pseudo-typed (PT) virus but cannot neutralize SARS-CoV or SARS-CoV-2 PT viruses [45, 46]. In contrast, 1249A8 neutralizes all three of the live viruses [33]. Both 1249A8 and B6 bind in the nanomolar range to SARS-CoV and SARS-CoV-2 S proteins, suggesting low S binding affinity is not responsible for B6’s inability to neutralize B-lineage viruses. In addition, SARS-CoV-2 PT virus neutralization assays were performed with B6 concentrations as high as 1mg/mL [45], a concentration >6,500x higher than B6’s *K*D for B-lineage virus S proteins, yet no neutralizing activity was observed [45].

How does one reconcile the apparent high NAb binding affinity for S proteins and peptides *in vitro*, with the inability to neutralize select CoV? We suggest that despite generally similar membrane fusion strategies, CoV S protein fusion mechanisms are unique amongst different CoV lineages. As a result, different viruses are more, or less, sensitive to different NAb neutralization mechanisms. For example, MERS-CoV is inhibited by C1 human (1249A8) and murine (B6, Fab22) NAbs that exhibit binding specificity for the MERS-CoV SH. These three NAbs all sterically disrupt 6HB1 formation and bind in similar orientations on the SH. However, analysis of the murine B6-SHp structures shows B6 does not lock the SARS-CoV-2 SH into a pre-fusion SH conformation. Thus, B6 is not expected to disrupt SH secondary structural changes required for SARS-CoV-2 S to transition from its PreF to PostF conformation, providing a possible explanation for B6’s inability to block SARS-CoV-2 PT viral infections. Thus, 1249A8 is unique in its ability to neutralize MERS-CoV, SARS-CoV, and SARS-CoV-2 through both steric and secondary structural disruption of S required for membrane fusion.

Although the SHp is small linear epitope, our study suggests SH-targeting NAbs may neutralize distinct CoV lineages using different molecular mechanisms. This idea has implications for immunization strategies that seek to induce BNAbs against multiple CoV. At least in mice, immunization with MERS-CoV S and SARS-CoV-2 S did not result in NAbs that neutralize both viruses [45, 46]. It remains unclear if this was due to the murine immune repertoire, the S antigens used in the vaccination experiments, the dosing regimens, or other unknown factors.

In humans, the history of endemic viral infections and their relationship to BNAb generation remains poorly characterized. Overall, our work suggests additional studies are required to define optimal vaccine strategies that induce BNMAbs against different CoV lineages, including those targeted by 1249A8.

## Methods

### Protein Expression and Fab generation

1249A8 NAb was expressed in Expi293 cells (Thermo Fisher, GIBCO) using manufacturer’s instructions and purified from the culture media using protein A affinity chromatography using MAbs select protein A resin (Cytiva). Fab was generated by papain (Sigma) digestion of 1249A8. Following digestion, the Fc was removed from the reaction using protein A resin. 1249A8 Fab was further purified by gel filtration chromatography. Purified 1249A8 Fab was incubated with a 5x molar excess of MERS-CoV SHp residues 1223-1245 for 30 minutes and the complex was concentrated to 10mg/mL for crystallization.

### SPR

SPR experiments were performed on a Biacore T200 (Cytiva) at 25°C using a running buffer consisting of 10mM HEPES, 150mM NaCl, 0.0075% P20. Binding studies were performed by capturing the hmAbs to the chip surface of CM-5 chips using a human antibody capture kit (cytiva). Kinetic binding parameters were derived by SPR using previously described SH-FC consisting of SH residues 1131-1171 [33]. SH-FC was injected over NAbs at four concentrations (25 nM, 6.125 nM, 1.56 nM, and 0.391 nM) with a contact time of 240 seconds and a 300 second dissociation time. All SPR experiments were double referenced (e.g., sensorgram data was subtracted from a control surface and from a buffer blank injection). The control surface for all experiments consisted of the capture antibody. Sensorgrams were globally fit to a 1:1 model, without a bulk shift correction, using Biacore T-200 evaluation software version 1.0. Fractional binding of SH-FC and CoV S proteins to NAbs was performed using the same injection parameters as for the kinetic experiments. Experiments were repeated at 25C, 30C, and 37C and due to minimal differences, analyzed as 3 replicates. S binding levels in RU, were converted to a fractional Rmax value by dividing RUobs by RUmax, where RUmax = NAb captured (RU) * [MW of S or SH-FC / 150,000], where the MW of all trimeric S proteins was fixed at 525,000 and the MW of SH-FC was 54,000. S proteins used in the study were SARS-CoV-2 S GCN4-IZ-2P (10561-CV, R&D Systems), SARS-CoV-2 B.1.1.529 S GCN4-IZ-2P (11061-CV, R&D Systems), MERS-CoV S GCN4-IZ-2P (R&D Systems), and SARS-CoV-2 B1.1.529 S (40589-V08H26, Sino Biological).

### Crystallization and Structure Determination

Crystals of the 1249A8 Fab / MERS-CoV SHp complex (in 20 mM NaPO4, pH 7,4, 100 mM NaCL, 10 mg/mL), were obtained from sitting drop vapor diffusion experiments, performed at 20C. All crystallization screens were performed using a Mosquito robot (SPT Labtech). Crystals were obtained in 200 nL drops consisting of 100 nL of complex with 100 nL well solution consisting of 0.2 M MgCl_2_·6H_2_O, 0.1M Tris, pH 8.5, 20% PEG-8000. Crystals, measuring 50 μm in each direction, were flash frozen in 25% glycerol 150 mM MgCl_2_·6H_2_O, 75 mM Tris, pH 8.5, 15% PEG-8000. Data were collected at the SER-CAT beamline 22-ID, Argonne National lab. The data were indexed, integrated, and scaled at 2.1Å resolution using DIALS [50], in the CCP4 package [51]. The structure was solved by molecular replacement using a model of 1249A8 Fab in Phenix. Model building was initially performed with ArpWarp [52, 53] and final modeling building and refinement was performed using Coot [54] and Phenix [55], respectively.

## Data Availability

Data and materials used and/or analyzed in this study are available from the corresponding author on reasonable request.

## Acknowledgements

We thank Sarah Moss for assistance with antibody purification. Use of the Biacore T-200 was made possible by the University of Alabama at Birmingham Multidisciplinary Molecular Interaction Core Facility. Data were collected at Southeast Regional Collaborative Access Team (SER-CAT) 22-ID beamline.

## References

1. Walsh EE, Shin JH, Falsey AR. Clinical impact of human coronaviruses 229E and OC43 infection in diverse adult populations. The Journal of infectious diseases. 2013;208(10):1634–42. Epub 2013/08/08. doi: 10.1093/infdis/jit393. PubMed PMID: 23922367; PubMed Central PMCID: PMCPMC3805243.

2. Lau SK, Woo PC, Yip CC, Tse H, Tsoi HW, Cheng VC, et al. Coronavirus HKU1 and other coronavirus infections in Hong Kong. Journal of clinical microbiology. 2006;44(6):2063–71. Epub 2006/06/08. doi: 10.1128/jcm.02614-05. PubMed PMID: 16757599; PubMed Central PMCID: PMCPMC1489438.

3. Wevers BA, van der Hoek L. Recently discovered human coronaviruses. Clinics in laboratory medicine. 2009;29(4):715–24. Epub 2009/11/07. doi: 10.1016/j.cll.2009.07.007. PubMed PMID: 19892230; PubMed Central PMCID: PMCPMC7131583.

4. Ksiazek TG, Erdman D, Goldsmith CS, Zaki SR, Peret T, Emery S, et al. A novel coronavirus associated with severe acute respiratory syndrome. The New England journal of medicine. 2003;348(20):1953–66. Epub 2003/04/12. doi: 10.1056/NEJMoa030781. PubMed PMID: 12690092.

5. Ksiazek TG, Erdman D, Goldsmith CS, Zaki SR, Peret T, Emery S, et al. Available from: https://www.cdc.gov/sars/about/fs-sars.html.

6. WHO MERS Global Summary and Assessment of Risk. 2018.

7. WHO:Home/Disease Outbreak News/Item/Middle East respiratory syndrome coronavirus (MERS-CoV) – Qatar. 2022.

8. Zhu N, Zhang D, Wang W, Li X, Yang B, Song J, et al. A Novel Coronavirus from Patients with Pneumonia in China, 2019. The New England journal of medicine. 2020;382(8):727–33. Epub 2020/01/25. doi: 10.1056/NEJMoa2001017. PubMed PMID: 31978945; PubMed Central PMCID: PMCPMC7092803.

9. WHO COVID-19 Dashboard: https://covid19.who.int/.

10. Woo PC, Lau SK, Lam CS, Lau CC, Tsang AK, Lau JH, et al. Discovery of seven novel Mammalian and avian coronaviruses in the genus deltacoronavirus supports bat coronaviruses as the gene source of alphacoronavirus and betacoronavirus and avian coronaviruses as the gene source of gammacoronavirus and deltacoronavirus. Journal of virology. 2012;86(7):3995–4008. Epub 2012/01/27. doi: 10.1128/jvi.06540-11. PubMed PMID: 22278237; PubMed Central PMCID: PMCPMC3302495.

11. Su S, Wong G, Shi W, Liu J, Lai ACK, Zhou J, et al. Epidemiology, Genetic Recombination, and Pathogenesis of Coronaviruses. Trends in microbiology. 2016;24(6):490–502. Epub 2016/03/26. doi: 10.1016/j.tim.2016.03.003. PubMed PMID: 27012512; PubMed Central PMCID: PMCPMC7125511.

12. Li W, Shi Z, Yu M, Ren W, Smith C, Epstein JH, et al. Bats are natural reservoirs of SARS-like coronaviruses. Science (New York, NY). 2005;310(5748):676–9. Epub 2005/10/01. doi: 10.1126/science.1118391. PubMed PMID: 16195424.

13. Memish ZA, Mishra N, Olival KJ, Fagbo SF, Kapoor V, Epstein JH, et al. Middle East respiratory syndrome coronavirus in bats, Saudi Arabia. Emerging infectious diseases. 2013;19(11):1819–23. Epub 2013/11/12. doi: 10.3201/eid1911.131172. PubMed PMID: 24206838; PubMed Central PMCID: PMCPMC3837665.

14. Anthony SJ, Gilardi K, Menachery VD, Goldstein T, Ssebide B, Mbabazi R, et al. Further Evidence for Bats as the Evolutionary Source of Middle East Respiratory Syndrome Coronavirus. mBio. 2017;8(2). Epub 2017/04/06. doi: 10.1128/mBio.00373-17. PubMed PMID: 28377531; PubMed Central PMCID: PMCPMC5380844.

15. Ruiz-Aravena M, McKee C, Gamble A, Lunn T, Morris A, Snedden CE, et al. Ecology, evolution and spillover of coronaviruses from bats. Nature reviews Microbiology. 2022;20(5):299–314. Epub 2021/11/21. doi: 10.1038/s41579-021-00652-2. PubMed PMID: 34799704; PubMed Central PMCID: PMCPMC8603903.

16. Cui J, Li F, Shi ZL. Origin and evolution of pathogenic coronaviruses. Nature reviews Microbiology. 2019;17(3):181–92. Epub 2018/12/12. doi: 10.1038/s41579-018-0118-9. PubMed PMID: 30531947; PubMed Central PMCID: PMCPMC7097006.

17. Lau SK, Li KS, Huang Y, Shek CT, Tse H, Wang M, et al. Ecoepidemiology and complete genome comparison of different strains of severe acute respiratory syndrome-related Rhinolophus bat coronavirus in China reveal bats as a reservoir for acute, self-limiting infection that allows recombination events. Journal of virology. 2010;84(6):2808–19. Epub 2010/01/15. doi: 10.1128/jvi.02219-09. PubMed PMID: 20071579; PubMed Central PMCID: PMCPMC2826035.

18. Hu B, Zeng LP, Yang XL, Ge XY, Zhang W, Li B, et al. Discovery of a rich gene pool of bat SARS-related coronaviruses provides new insights into the origin of SARS coronavirus. PLoS pathogens. 2017;13(11):e1006698. Epub 2017/12/01. doi: 10.1371/journal.ppat.1006698. PubMed PMID: 29190287; PubMed Central PMCID: PMCPMC5708621.

19. Jackson B, Boni MF, Bull MJ, Colleran A, Colquhoun RM, Darby AC, et al. Generation and transmission of interlineage recombinants in the SARS-CoV-2 pandemic. Cell. 2021;184(20):5179–88.e8. Epub 2021/09/10. doi: 10.1016/j.cell.2021.08.014. PubMed PMID: 34499854; PubMed Central PMCID: PMCPMC8367733.

20. Davies NG, Abbott S, Barnard RC, Jarvis CI, Kucharski AJ, Munday JD, et al. Estimated transmissibility and impact of SARS-CoV-2 lineage B.1.1.7 in England. Science (New York, NY). 2021;372(6538):eabg3055. doi: 10.1126/science.abg3055.

21. Menachery VD, Yount BL, Jr., Debbink K, Agnihothram S, Gralinski LE, Plante JA, et al. A SARS-like cluster of circulating bat coronaviruses shows potential for human emergence. Nat Med. 2015;21(12):1508–13. Epub 2015/11/10. doi: 10.1038/nm.3985. PubMed PMID: 26552008; PubMed Central PMCID: PMCPMC4797993.

22. Li H, Mendelsohn E, Zong C, Zhang W, Hagan E, Wang N, et al. Human-animal interactions and bat coronavirus spillover potential among rural residents in Southern China. Biosafety and health. 2019;1(2):84–90. Epub 2020/06/06. doi: 10.1016/j.bsheal.2019.10.004. PubMed PMID: 32501444; PubMed Central PMCID: PMCPMC7148670.

23. Wrapp D, Wang N, Corbett KS, Goldsmith JA, Hsieh C-L, Abiona O, et al. Cryo-EM structure of the 2019-nCoV spike in the prefusion conformation. Science (New York, NY). 2020;367(6483):1260–3. doi: 10.1126/science.abb2507.

24. Walls AC, Park YJ, Tortorici MA, Wall A, McGuire AT, Veesler D. Structure, Function, and Antigenicity of the SARS-CoV-2 Spike Glycoprotein. Cell. 2020;181(2):281–92 e6. Epub 2020/03/11. doi: 10.1016/j.cell.2020.02.058. PubMed PMID: 32155444; PubMed Central PMCID: PMCPMC7102599.

25. Yuan Y, Cao D, Zhang Y, Ma J, Qi J, Wang Q, et al. Cryo-EM structures of MERS-CoV and SARS-CoV spike glycoproteins reveal the dynamic receptor binding domains. Nature communications. 2017;8:15092. Epub 2017/04/11. doi: 10.1038/ncomms15092. PubMed PMID: 28393837; PubMed Central PMCID: PMCPMC5394239.

26. Cai Y, Zhang J, Xiao T, Peng H, Sterling SM, Walsh RM, et al. Distinct conformational states of SARS-CoV-2 spike protein. Science (New York, NY). 2020;369(6511):1586–92. doi: doi:10.1126/science.abd4251.

27. Wang N, Shi X, Jiang L, Zhang S, Wang D, Tong P, et al. Structure of MERS-CoV spike receptor-binding domain complexed with human receptor DPP4. Cell Res. 2013;23(8):986–93. doi: 10.1038/cr.2013.92. PubMed PMID: 23835475.

28. Liu H, Wei P, Kappler JW, Marrack P, Zhang G. SARS-CoV-2 Variants of Concern and Variants of Interest Receptor Binding Domain Mutations and Virus Infectivity. Frontiers in immunology. 2022;13. doi: 10.3389/fimmu.2022.825256.

29. Piccoli L, Park YJ, Tortorici MA, Czudnochowski N, Walls AC, Beltramello M, et al. Mapping Neutralizing and Immunodominant Sites on the SARS-CoV-2 Spike Receptor-Binding Domain by Structure-Guided High-Resolution Serology. Cell. 2020;183(4):1024–42.e21. Epub 2020/09/30. doi: 10.1016/j.cell.2020.09.037. PubMed PMID: 32991844; PubMed Central PMCID: PMCPMC7494283.

30. Wec AZ, Wrapp D, Herbert AS, Maurer D, Haslwanter D, Sakharkar M, et al. Broad sarbecovirus neutralizing antibodies define a key site of vulnerability on the SARS-CoV-2 spike protein. bioRxiv. 2020. Epub 2020/06/09. doi: 10.1101/2020.05.15.096511. PubMed PMID: 32511337; PubMed Central PMCID: PMCPMC7241100.

31. McLean G, Kamil J, Lee B, Moore P, Schulz TF, Muik A, et al. The Impact of Evolving SARS-CoV-2 Mutations and Variants on COVID-19 Vaccines. mBio. 2022;13(2):e0297921. Epub 2022/03/31. doi: 10.1128/mbio.02979-21. PubMed PMID: 35352979; PubMed Central PMCID: PMCPMC9040821.

32. Tada T, Zhou H, Dcosta BM, Samanovic MI, Chivukula V, Herati RS, et al. Increased resistance of SARS-CoV-2 Omicron variant to neutralization by vaccine-elicited and therapeutic antibodies. EBioMedicine. 2022;78:103944. Epub 2022/04/26. doi: 10.1016/j.ebiom.2022.103944. PubMed PMID: 35465948; PubMed Central PMCID: PMCPMC9021600.

33. Piepenbrink MS, Park JG, Desphande A, Loos A, Ye C, Basu M, et al. Potent universal-coronavirus therapeutic activity mediated by direct respiratory administration of a Spike S2 domain-specific human neutralizing monoclonal antibody. bioRxiv. 2022. Epub 2022/03/17. doi: 10.1101/2022.03.05.483133. PubMed PMID: 35291292; PubMed Central PMCID: PMCPMC8923109.

34. Hurlburt NK, Homad LJ, Sinha I, Jennewein MF, MacCamy AJ, Wan YH, et al. Structural definition of a pan-sarbecovirus neutralizing epitope on the spike S2 subunit. Communications biology. 2022;5(1):342. Epub 2022/04/13. doi: 10.1038/s42003-022-03262-7. PubMed PMID: 35411021; PubMed Central PMCID: PMCPMC9001700 63/131599 filed by the Fred Hutchinson Cancer Research Center on the CV3-25 monoclonal antibody. A.T.M. is an inventor on a provisional patent application No. 63/108,554 filed by the University of Washington on the B6 monoclonal antibody. All other authors declare no competing interests.

35. Pinto D, Sauer MM, Czudnochowski N, Low JS, Tortorici MA, Housley MP, et al. Broad betacoronavirus neutralization by a stem helix-specific human antibody. Science (New York, NY). 2021;373(6559):1109–16. Epub 2021/08/05. doi: 10.1126/science.abj3321. PubMed PMID: 34344823.

36. Zhou P, Yuan M, Song G, Beutler N, Shaabani N, Huang D, et al. A human antibody reveals a conserved site on beta-coronavirus spike proteins and confers protection against SARS-CoV-2 infection. Science translational medicine. 2022;14(637):eabi9215. Epub 2022/02/09. doi: 10.1126/scitranslmed.abi9215. PubMed PMID: 35133175; PubMed Central PMCID: PMCPMC8939767.

37. Li W, Chen Y, Prévost J, Ullah I, Lu M, Gong SY, et al. Structural basis and mode of action for two broadly neutralizing antibodies against SARS-CoV-2 emerging variants of concern. Cell reports. 2022;38(2):110210. Epub 2022/01/01. doi: 10.1016/j.celrep.2021.110210. PubMed PMID: 34971573; PubMed Central PMCID: PMCPMC8673750.

38. Jennewein MF, MacCamy AJ, Akins NR, Feng J, Homad LJ, Hurlburt NK, et al. Isolation and characterization of cross-neutralizing coronavirus antibodies from COVID-19+ subjects. Cell reports. 2021;36(2):109353. Epub 2021/07/09. doi: 10.1016/j.celrep.2021.109353. PubMed PMID: 34237283; PubMed Central PMCID: PMCPMC8216847.

39. Adams MD, Celniker SE, Holt RA, Evans CA, Gocayne JD, Amanatides PG, et al. The genome sequence of Drosophila melanogaster. Science (New York, NY). 2000;287(5461):2185–95. PubMed PMID: 10731132.

40. Pallesen J, Wang N, Corbett KS, Wrapp D, Kirchdoerfer RN, Turner HL, et al. Immunogenicity and structures of a rationally designed prefusion MERS-CoV spike antigen. Proceedings of the National Academy of Sciences of the United States of America. 2017;114(35):E7348–e57. Epub 2017/08/16. doi: 10.1073/pnas.1707304114. PubMed PMID: 28807998; PubMed Central PMCID: PMCPMC5584442.

41. Kirchdoerfer RN, Cottrell CA, Wang N, Pallesen J, Yassine HM, Turner HL, et al. Pre-fusion structure of a human coronavirus spike protein. Nature. 2016;531(7592):118–21. Epub 2016/03/05. doi: 10.1038/nature17200. PubMed PMID: 26935699; PubMed Central PMCID: PMCPMC4860016.

42. Turoňová B, Sikora M, Schürmann C, Hagen WJH, Welsch S, Blanc FEC, et al. In situ structural analysis of SARS-CoV-2 spike reveals flexibility mediated by three hinges. Science (New York, NY). 2020;370(6513):203–8. Epub 2020/08/21. doi: 10.1126/science.abd5223. PubMed PMID: 32817270; PubMed Central PMCID: PMCPMC7665311.

43. Ke Z, Oton J, Qu K, Cortese M, Zila V, McKeane L, et al. Structures and distributions of SARS-CoV-2 spike proteins on intact virions. Nature. 2020;588(7838):498–502. Epub 2020/08/18. doi: 10.1038/s41586-020-2665-2. PubMed PMID: 32805734; PubMed Central PMCID: PMCPMC7116492.

44. Hsieh CL, Goldsmith JA, Schaub JM, DiVenere AM, Kuo HC, Javanmardi K, et al. Structure-based design of prefusion-stabilized SARS-CoV-2 spikes. Science (New York, NY). 2020;369(6510):1501–5. Epub 2020/07/25. doi: 10.1126/science.abd0826. PubMed PMID: 32703906; PubMed Central PMCID: PMCPMC7402631.

45. Sauer MM, Tortorici MA, Park YJ, Walls AC, Homad L, Acton OJ, et al. Structural basis for broad coronavirus neutralization. Nature structural & molecular biology. 2021;28(6):478–86. Epub 2021/05/14. doi: 10.1038/s41594-021-00596-4. PubMed PMID: 33981021.

46. Hsieh CL, Werner AP, Leist SR, Stevens LJ, Falconer E, Goldsmith JA, et al. Stabilized coronavirus spike stem elicits a broadly protective antibody. Cell reports. 2021;37(5):109929. Epub 2021/10/29. doi: 10.1016/j.celrep.2021.109929. PubMed PMID: 34710354; PubMed Central PMCID: PMCPMC8519809.

47. Wang C, van Haperen R, Gutiérrez-Álvarez J, Li W, Okba NMA, Albulescu I, et al. A conserved immunogenic and vulnerable site on the coronavirus spike protein delineated by cross-reactive monoclonal antibodies. Nature communications. 2021;12(1):1715. Epub 2021/03/19. doi: 10.1038/s41467-021-21968-w. PubMed PMID: 33731724; PubMed Central PMCID: PMCPMC7969777 B.J.B. are inventors on a patent application on monoclonal antibodies targeting MERS-CoV (patent publication no.: WO/2020/169755). F.G., D.D. and R.H. are non-substantial interest shareholders in Harbour Biomed and were part of the team that generated the mice. The other authors declare no competing interests.

48. Li Y, Lai DY, Zhang HN, Jiang HW, Tian X, Ma ML, et al. Linear epitopes of SARS-CoV-2 spike protein elicit neutralizing antibodies in COVID-19 patients. Cellular & molecular immunology. 2020;17(10):1095–7. Epub 2020/09/09. doi: 10.1038/s41423-020-00523-5. PubMed PMID: 32895485; PubMed Central PMCID: PMCPMC7475724.

49. Polyiam K, Phoolcharoen W, Butkhot N, Srisaowakarn C, Thitithanyanont A, Auewarakul P, et al. Immunodominant linear B cell epitopes in the spike and membrane proteins of SARS-CoV-2 identified by immunoinformatics prediction and immunoassay. Scientific reports. 2021;11(1):20383. Epub 2021/10/16. doi: 10.1038/s41598-021-99642-w. PubMed PMID: 34650130; PubMed Central PMCID: PMCPMC8516869.

50. Winter G, Waterman DG, Parkhurst JM, Brewster AS, Gildea RJ, Gerstel M, et al. DIALS: implementation and evaluation of a new integration package. Acta crystallographica Section D, Structural biology. 2018;74(Pt 2):85–97. Epub 2018/03/14. doi: 10.1107/s2059798317017235. PubMed PMID: 29533234; PubMed Central PMCID: PMCPMC5947772.

51. Winn MD, Ashton AW, Briggs PJ, Ballard CC, Patel P. Ongoing developments in CCP4 for high-throughput structure determination. Acta crystallographica Section D, Biological crystallography. 2002;58(Pt 11):1929–36. PubMed PMID: 12393924.

52. Perrakis A, Sixma TK, Wilson KS, Lamzin VS. wARP: improvement and extension of crystallographic phases by weighted averaging of multiple-refined dummy atomic models. Acta crystallographica Section D, Biological crystallography. 1997;53(Pt 4):448–55. PubMed PMID: 15299911.

53. Perrakis A, Harkiolaki M, Wilson KS, Lamzin VS. ARP/wARP and molecular replacement. Acta crystallographica Section D, Biological crystallography. 2001;57(Pt 10):1445–50. Epub 2001/09/22. doi: 10.1107/s0907444901014007. PubMed PMID: 11567158.

54. Emsley P, Lohkamp B, Scott WG, Cowtan K. Features and development of Coot. Acta crystallographica Section D, Biological crystallography. 66(Pt 4):486–501. Epub 2010/04/13. doi: S0907444910007493 [pii] 10.1107/S0907444910007493. PubMed PMID: 20383002.

55. Adams PD, Afonine PV, Bunkoczi G, Chen VB, Echols N, Headd JJ, et al. The Phenix software for automated determination of macromolecular structures. Methods. 2011. Epub 2011/08/09. doi: S1046-2023(11)00131-9 [pii] 10.1016/j.ymeth.2011.07.005. PubMed PMID: 21821126.

