## Supplemental Table 1 and Fig. 1 for "Structure and Epitope of a Neutralizing Monoclonal Antibody that Targets the Stem Helix of β Coronaviruses"

**Supplemental Table 1. Data Collection**

| | |
|---|---|
| Beamline | SER-CAT 22-ID |
| Wavelength, Å | 1.0 |
| Space group | $P2_1$ |
| Unit cell parameters | |
| a, b, c, β | 52.714, 56.308, 76.124, 91.191 |
| Resolution, Å | 45.3 - 2.1 |
| Number of Obs. | 86,813 |
| Unique reflections | 26,008 |
| Redundancy[a] | 3.3 (3.0) |
| Completeness[a] | 98.6 (93.9) |
| <I/sigI>[a] | 5.0 (1.9) |
| Rmrg (%)[a] | 0.119 (0.436) |
| Rpim, (%)[a] | 0.082 (0.351) |
| CC1/2, (%)[a] | 0.967 (93.9) |

**Refinement Statistics**

| | |
|---|---|
| Resolution (Å) | 43.8 - 2.1 |
| Reflections (work) | 25,899 |
| Reflections (test) | 1,410 |
| Rcryst / Rfree | 18.17 / 23.84 |
| No. of atoms | |
| Protein | 3,433 |
| Solvent | 282 |
| Ions ($Mg^{+2}$, $Cl^-$) | 11 |
| Wilson B factor ($Å^2$) | 29.5 |
| Mean B factor ($Å^2$) | 31.0 |

**Model Geometry**

| | |
|---|---|
| Bond lengths (Å) | 0.0087 |
| Bond angles (°) | 1.02 |
| Dihedrals (°) | 17.875 |
| Ramachandran plot | |
| Favored (%) | 98.63 |
| Outliers (%) | 0.23 |
| PDB ID | processing submission to PDB |

[a] Numbers in parentheses refer to highest resolution shell (2.16-2.10Å).
CC1/2 = Pearson correlation coefficient between two random half datasets.

$$R_{\mathrm{mrg}} = \sum_{hkl}\sum_{i} |I_i(hkl) - \overline{I(hkl)}| \Big/ \sum_{hkl}\sum_{i} I_i(hkl) \quad .$$

$$R_{\mathrm{pim}} = \sum_{hkl}[1/(N-1)]^{1/2} \sum_{i} |I_i(hkl) - \overline{I(hkl)}| \Big/ \sum_{hkl}\sum_{i} I_i(hkl)$$

**Supplemental Figure 1.**

| | 2nd Struc. | | Res. | Sequence | | Buried Surface | | | | # Hydrogen Bonds | | | |
|---|---|---|---|---|---|---|---|---|---|---|---|---|---|
| | POST | PRE | # | MERS-CoV | SARS-CoV-2 | 1249A8 | S2P6 | CC40.8 | CV3-25 | 1249A8 | S2P6 | CC40.8 | CV3-25 |
| | - | - | 1223 | L | P | | | 0 | | | | | |
| | C | H | 1224 | G | L | | | 0.37 | | | | | |
| | C | H | 1225 | N | Q | | | 84.46 | | | | 2H | |
| | C | H | 1226 | S | P | | | 46.61 | | | | H | |
| | C | H | 1227 | T | E | | | 59.3 | | | | H | |
| | C | H | 1228 | G | L | | | 156.73 | | | | H | |
| | C | H | 1229 | I | D | | 4.34 | 92.32 | 0 | | | 3H | |
| | C | H | 1230 | D | S | 35.42 | 16.01 | 0 | 0 | | | | |
| CORE SH | **H** | **H** | 1231 | **F** | **F** | 122.89 | 162.44 | 67.81 | 0 | H | H | | |
| | **H** | **H** | 1232 | Q | K | 106.44 | 84.06 | 72.27 | 14.94 | 2H | H | | H |
| | **H** | **H** | 1233 | D | E | 0 | 0 | 12.55 | 10.79 | | | | |
| | **H** | **H** | 1234 | **E** | **E** | 56.78 | 26.77 | 65.71 | 0 | 2H | | H | |
| | **H** | **H** | 1235 | **L** | **L** | 81.58 | 74.81 | 109.17 | 23.69 | | | | |
| | **H** | **H** | 1236 | **D** | **D** | 18.72 | 13.48 | 51 | 86.28 | H | H | H | 4H |
| | **H** | **H** | 1237 | E | K | 0 | 0 | 43.95 | 5.18 | | | H | |
| | C | H | 1238 | F | Y | 73.71 | 73.58 | 102.17 | 0 | | | | |
| | C | H | 1239 | **F** | **F** | 118.6 | 122.66 | 73.77 | 44.4 | | | | H |
| | C | H | 1240 | **K** | **K** | 0 | 0 | 0 | 135.55 | | | | 5H |
| | C | H | 1241 | N | N | 0 | 0 | 4.91 | 53.2 | | | | |
| | C | H | 1242 | V | H | | 7.86 | 37.8 | 14.41 | | | | H |
| | C | H | 1243 | S | T | | | 16.06 | 56.85 | | | | 2H |
| | C | H | 1244 | T | S | | | 71.76 | 50.01 | | | 2H | 3H |
| | C | H | 1245 | S | P | | | | 34.47 | | | | |
| | C | - | 1246 | I | D | | | | 43.4 | | | | 3H |
| | C | - | 1247 | P | V | | | | 73.07 | | | | |
| | C | - | 1248 | N | D | | | | 10.5 | | | | |
| | | | | | **Totals >** | 614.14 | 586.01 | 1168.7 | 656.74 | 6 | 3 | 13 | 20 |
| | | | | | **CORE >** | 386.41 | 361.56 | 422.46 | 140.88 | 6 | 3 | 3 | 11 |
| | | | | | **#res >** | 12 | 14 | 22 | 20 | | | | |
